# Genomic and epidemiological evidence of a dominant Panton-Valentine leucocidin-positive Methicillin Resistant *Staphylococcus aureus* lineage in Sri Lanka with spread to the United Kingdom and Australia

**DOI:** 10.1101/521260

**Authors:** S.M. McTavish, S.J. Snow, E.C. Cook, B. Pichon, S. Coleman, G.W. Coombs, S. Pang, C.A. Arias, L. Díaz, E. Boldock, S. Davies, M. Udukala, A. Kearns, S. Siribaddana, T.I. de Silva

**Author notes:** These authors contributed equally to this work (S.M. McTavish, S.J. Snow, E.C. Cook). Corresponding author: Thushan I. de Silva The Florey Institute for Host-Pathogen Interactions, Department of Infection, Immunity and Cardiovascular Disease, University of Sheffield, Beech Hill Road, Sheffield S10 2RX, UK.

## Abstract

**Objective:** To undertake the first detailed genomic analysis of methicillin-resistant *Staphylococcus aureus* (MRSA) isolated in Sri Lanka.

**Methods:** A prospective observational study was performed on 94 MRSA isolates collected over a four month period from the Anuradhapura Teaching Hospital, Sri Lanka. Screening for *mec*A, *mec*C and the Panton-Valentine leucocidin (PVL)-associated *lukS-PV/lukF-PV* genes and molecular characterisation by *spa* typing was undertaken. Whole genome sequencing (WGS) and phylogenetic analysis was performed on selected multilocus sequence type (MLST) clonal complex 5 (CC5) isolates from Sri Lanka, England, Australia and Argentina.

**Results:** All 94 MRSA harboured the *mecA* gene. Nineteen *spa* types associated with nine MLST clonal complexes were identified. Most isolates were from skin and soft tissue infections (76.9%), with the remainder causing more invasive disease. Sixty two (65.9%) of isolates were PVL positive with the majority (56 isolates; 90.3%) belonging to a dominant CC5 lineage. This lineage, PVL-positive ST5-MRSA-IVc, was associated with community and hospital-onset infections. Based on WGS, representative PVL-positive ST5-MRSA-IVc isolates from Sri Lanka, England and Australia formed a single phylogenetic clade, suggesting wide geographical circulation.

**Conclusions:** We present the most detailed genomic analysis of MRSA isolated in Sri Lanka to date. The analysis identified a PVL-positive ST5-MRSA-IVc that dominates MRSA clinical infections in Sri Lanka. Furthermore, transmission of the strain has occurred in the United Kingdom and Australia.

## Introduction

Worldwide, *Staphylococcus aureus* is the primary causative agent of community-acquired skin and soft tissue infections (SSTI). The emergence of methicillin-resistant *S. aureus* (MRSA) has led to *S. aureus* becoming an important cause of hospital-associated invasive infections including bacteraemia, pneumonia and endocarditis (Bell et al., 2002;David and Daum, 2010). Panton-Valentine leucocidin (PVL)-positive MRSA is a well-documented cause of community-associated SSTI and less commonly, life-threatening infections in immunocompetent populations. Its prevalence is thought to be increasing worldwide and multi drug resistant PVL-MRSA is emerging as a threat, particularly in the Indian subcontinent (Song et al., 2011;Shallcross et al., 2013). In many developed countries, surveillance of MRSA invasive disease, characterisation of high risk MRSA clones and the investigation of suspected MRSA outbreaks are achieved through public health tracking and molecular analysis. By comparison, limited data exist on MRSA infections in low and middle-income countries. A recent report has suggested Sri Lankan hospitals have the highest prevalence of MRSA for all Asian hospitals that were included in the study (Song et al., 2011). However, information on the molecular epidemiology and spectrum of clinical disease is lacking (Corea et al., 2003;Mahalingam et al., 2014;Jayaweera et al., 2017;Jayaweera and Kumbukgolla, 2017). Consequently in our study, we report on the genomic analysis of MRSA isolated from patients admitted to a major teaching hospital in Sri Lanka.

## Methods

A prospective, observational study of sequential MRSA infections in hospitalised patients was conducted at the Anuradhapura Teaching Hospital from 30^th^ June to 31^st^ October 2014. This hospital serves approximately 1.6 million people living in the rural North Central province of Sri Lanka. MRSA was identified from clinical specimens via disc diffusion using oxacillin and CLSI criteria. All MRSA isolated from any site with a clinical infection during the 4-month period were included in the study. Ethical approval was obtained from the Ethics Review Committee, Rajarata University of Sri Lanka. Infections were defined as community-onset (CO) if the sample was collected <72 hours from admission and hospital-onset (HO) if collected later, based on previous studies distinguishing whether MRSA infection were associated with community or hospital settings (Maree et al., 2007).

Isolates were referred to the Staphylococcal Reference Service, National Infection Service, Public Health England (PHE), Colindale, London for further analysis. Initial testing was performed using the MALDI-TOF (MALDI Biotyper^®^, Bruker Daltonik GmbH, Germany), followed by reverse-transcriptase polymerase chain reaction (RT-PCR) for *mecA*, *mecC* and *lukS*-PV/*lukF*-PV genes, to determine the isolate’s methicillin resistance, Panton-Valentine leucocidin (PVL) status, and *spa* typing (Frenay et al., 1996;Pichon et al., 2012). Whole genome sequencing (WGS) on selected isolates was undertaken as previously described (Garvey et al., 2016;Lahuerta-Marin et al., 2016). Phylogeny was inferred by maximum likelihood analysis using RAxML and the GTRCAT model.

## Results

The 94 isolates submitted for further testing were confirmed as *S. aureus* by MALDI-TOF and were *mec*A positive (Table 1). No samples were *mecC* positive. Where clinical data were available (n=91), the majority of MRSA isolates (n=70, 76.9%) were from skin and soft tissue infections (SSTIs), with the remainder from invasive infections, including 16 (17.6%) MRSA bacteraemias (Table 1). Based on the 19 *spa* types identified, the isolates could be grouped into nine MLST clonal complexes (CC) including: CC5 (n=59 isolates), CC30 (n=18), CC1 (n=8), CC59 (n=4), and single isolates belonging to CC6, CC8, CC45, CC97 and CC101. The dominant CC5 MRSA lineage (62.7% of isolates) was comprised of six related *spa* types: t002 (n=51), t010 (n=4) and single isolates of t045, t1062, t5490 and t7342. Sixty two (65.9%) isolates were PVL positive, the majority (56 isolates; 90.3%) belonging to the CC5 lineage. The CC5 PVL-positive lineage was associated mainly with HO- and CO-SSTIs (42; 84.0%), but was also responsible for more invasive infections (Table 1). All HO-SSTIs were surgical wound infections.

**Table 1.**
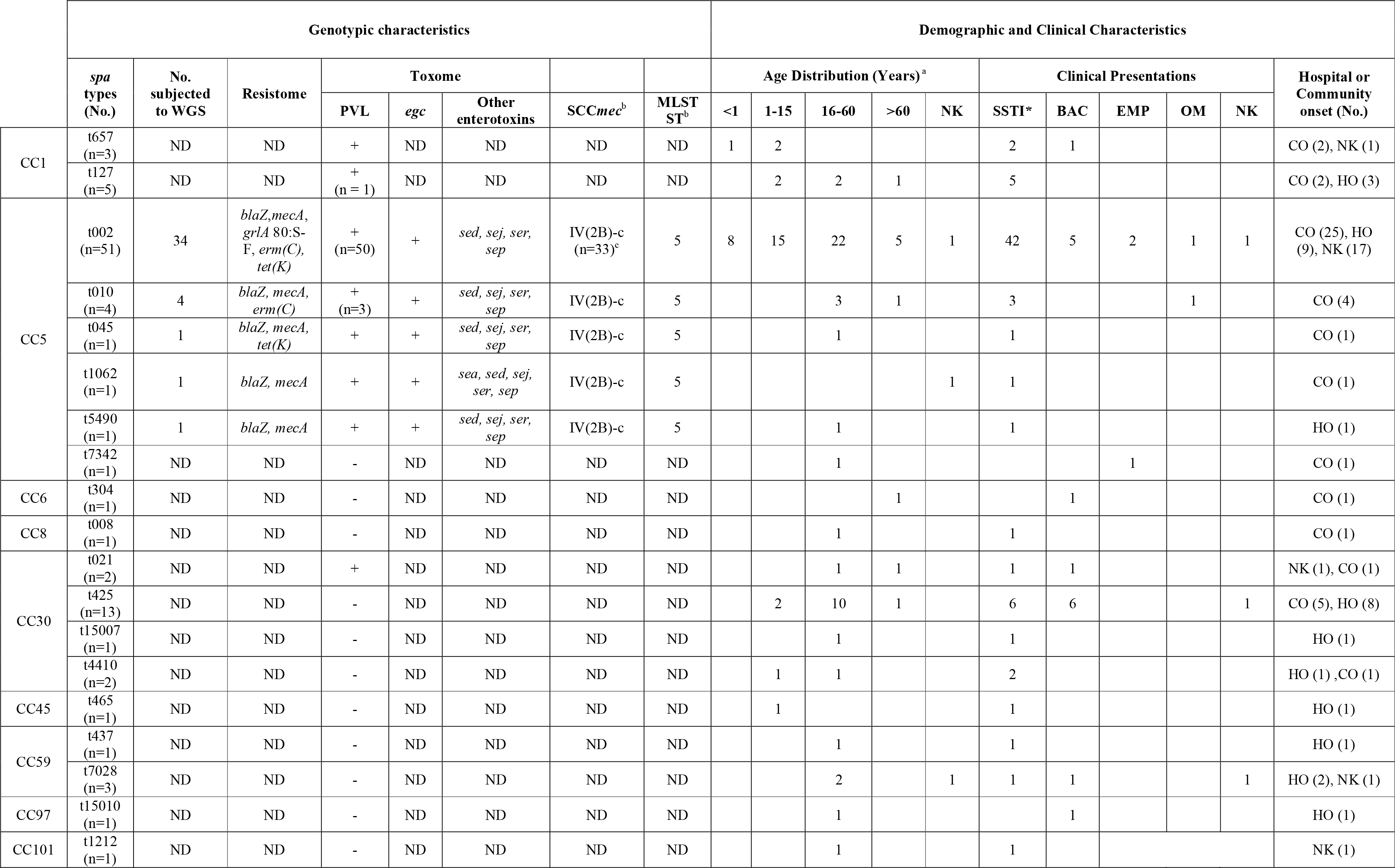
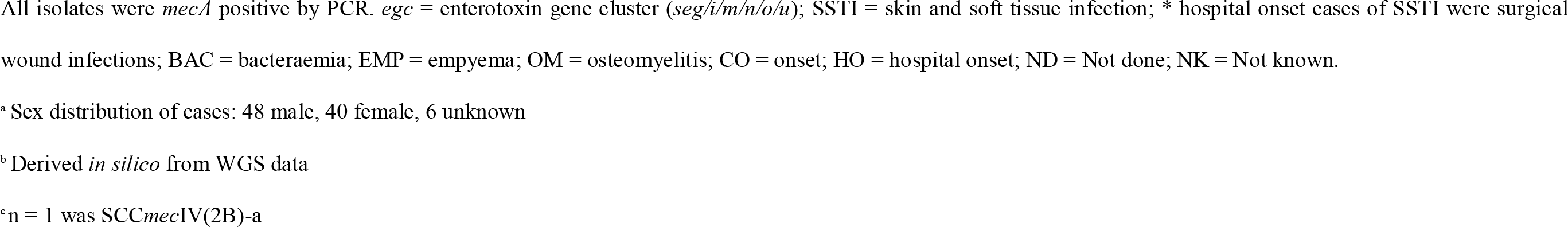
Genotypic, demographic and clinical characteristics of MRSA isolates from Sri Lanka.

To investigate whether the CC5 PVL-positive lineage was genotypically diverse, or a single circulating clone, WGS was performed on 41 isolates selected on the basis of diverse clinical symptoms and *spa* type (34 t002 and 7 samples from other *spa* types). As CC5 PVL-positive MRSA have been identified sporadically in England, we sought to assess the relatedness of the Sri Lankan isolates to lineage-matched isolates from held in the PHE archives. These included isolates from patients with known links to Sri Lanka (10 CC5 PVL-positive MRSA isolates (dates of collection between 2005 and 2014)) and patients with no known links to Sri Lanka (79 isolates: 12 CC5 PVL-positive MRSA (2005-2015), 33 CC5 PVL-negative MRSA (2009-2016), 4 CC5 PVL-positive methicillin-sensitive *S. aureus* (MSSA) (2011-2016) and 30 CC5 PVL-negative MSSA (2011-2016)). Previously sequenced CC5 PVL-positive MRSA from collaborators in Australia (n=14, collected in 2015) and Argentina (n=3; collected in 2003) were also included as comparators.

Phylogenetic analysis of CC5 strain WGS showed great variability (Figure 1A), with isolates from various countries dispersed throughout the tree. A strong geographic signal however was apparent amongst the isolates from Sri Lanka, with all but three CC5-PVL-positive MRSA isolates clustering into a single clade, herein dubbed the “Sri Lankan clade” (Figure 1B). Isolates from the United Kingdom (13) and Australia (1) were also found in the Sri Lankan clade, including those from patients with no known links to Sri Lanka. Within the clade, the isolates were identified as multilocus sequence type (ST) 5, harboured the enterotoxin gene cluster (*egc*), and for the MRSA isolates harboured the SCC*mec* IVc staphylococcal cassette chromosome *mec* subtype. All but one isolate encoded the plasmid-borne enterotoxin genes (*sed*, *sej* and *ser*). Greater variability was apparent for other traits including the *sep* enterotoxin gene. Genes encoding resistance to erythromycin (*erm*(C)) or tetracycline (*tet*(K)) were variably detected highlighting the dynamic loss/acquisition of mobile genetic elements within the clone. Similarly, a chromosomal mutation associated with quinolone resistance (*grlA* 80:S-F) was noted sporadically. A single isolate from a UK patient with links to Sri Lanka was identified as being genotypically multi-drug resistant, encoding *blaZ*, *mecA*, *erm*(C), *tet*(K), *aphA3* and *sat4* genes. Bayesian phylogenetic reconstruction using BEAST (data not shown) failed to provide significant temporal signal for predicting evolutionary rate and time to common ancestor.

**Figure 1.**
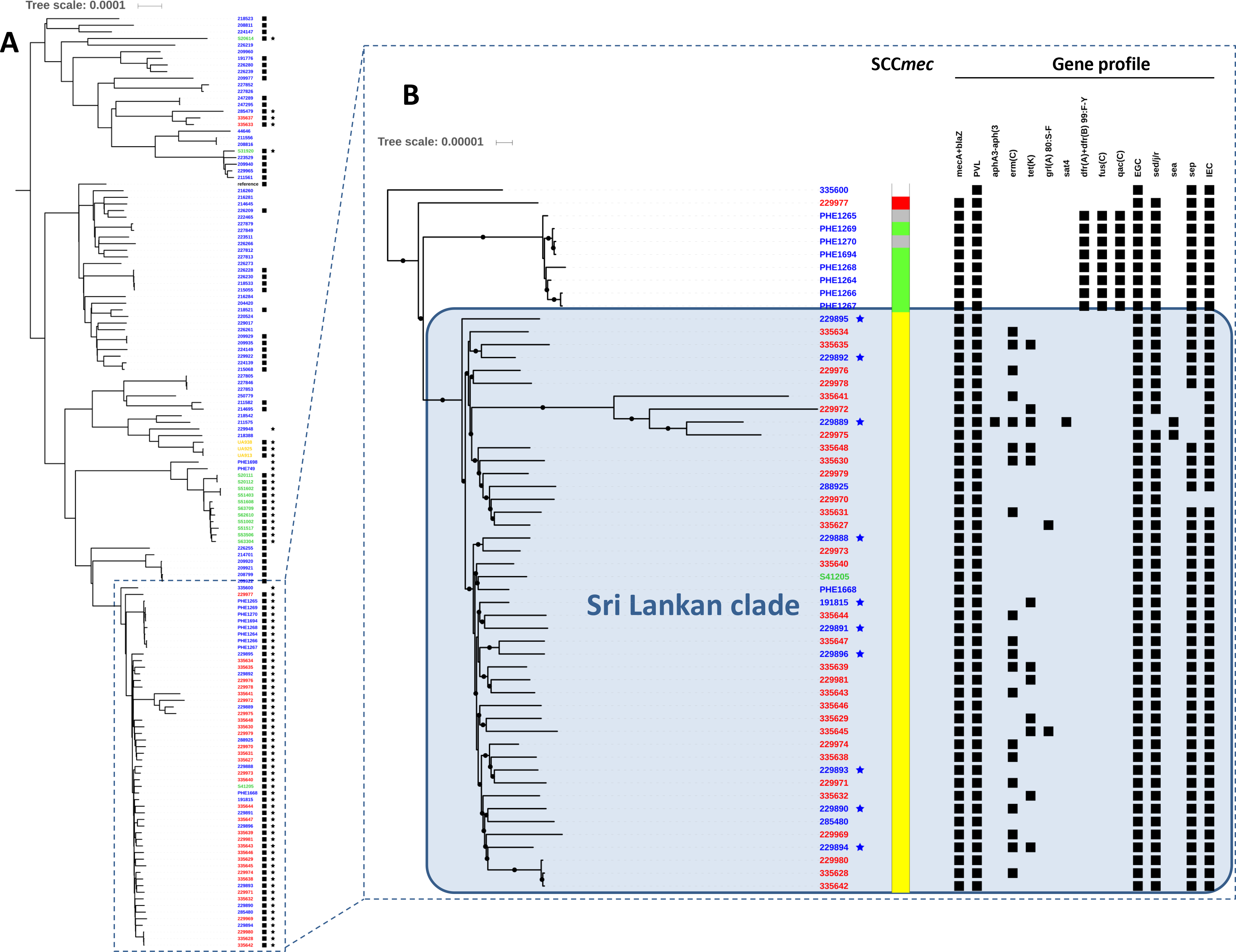
Alignment of international MLST CC5 *Staphylococcus aureus* genomes. **A).** Phylogenetic tree indicating relationships between international CC5 *S. aureus* including their PVL (★) and *mecA* (▀) status based on SNP analysis of whole genome sequences. Phylogeny was inferred by maximum likelihood analysis using RAxML GTRCAT model with 100 bootstraps from aligned polymorphic sites allowing 20% of Ns and gaps. Polymorphic sites were called using gatk2 and filtered (AD ratio =0.9; min depth =10; MQ score > 30; QUAL score >40) using genome NC_002745 as mapping reference. The tree was drawn using FigTree v1.4.3. Country of origin denoted as follows; blue: England; red: Sri Lanka; green: Australia; yellow: Argentina. Scale is in substitutions per site and indicates ≈ 130 SNPs. **B)** 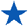 Indicates UK patients with known links to Sri Lanka. □ MSSA; SCC*mec* types: 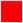 IV-a; 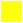: IV-c; 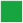 VI; 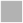 NT. In gene profile section, ▀ indicates presence of gene. Scale is in substitutions per site and represents ≈ 13 SNPs.

## Discussion

Particular lineages of MRSA are frequently associated with various geographical origins e.g. ST8/USA300 (North America); ST93 (Australia); ST80 (North Africa); ST30 (South-West Pacific) (David and Daum, 2010;Chua et al., 2011). Prior to the current study, CC5 PVL-positive MRSA has been reported in many countries world-wide (Monecke et al., 2011); however, their origin(s) are unclear. A recent phylogenomic study of CC5-MRSA isolates from the Western Hemisphere showed high diversity, even among strains that shared the same SCC*mec* type circulating in the same country (Challagundla et al., 2018). Very few whole genome sequences from CC5-MRSA isolates from Asian countries are currently available.

Herein, in most instances, phylogenetic analysis of CC5 PVL-positive MRSA from four continents showed clustering according to their geographic location, suggesting they have arisen independently in different parts of world following the acquisition of PVL phage and/or different SCC*mec* elements. Our data provide evidence of a successful ST5-PVL-positive MRSA-IVc clone in Sri Lanka with multiple incursions to distant geographical regions. Thirteen isolates from England were interspersed within the Sri Lankan clade; ten with known links to Sri Lanka. One isolate from Australia also clustered within the Sri Lankan clade, however a link to Sri Lanka could not be determined. Whilst in our study all Sri Lankan isolates were collected systematically without undue bias, it is important to acknowledge that the number of isolates is small and that they were collected from a single centre over a relatively short timeframe. However, the UK CC5 PVL-positive MRSA isolates in the Sri Lankan clade, including those with known links to Sri Lanka, were collected over the course of a decade prior to our study. This suggests that wider circulation of this PVL-positive ST5-MRSA-IVc clone is likely in Sri Lanka and that our newly collected samples do not simply represent a clonal outbreak in Anuradhapura Teaching Hospital. A larger study of isolates from other parts of Sri Lanka and globally is required to help elucidate the origins and dissemination of PVL-positive MRSA belonging to the CC5 lineage. The ST5-PVL-positive MRSA-IVc clone identified was also responsible for both CO- and HO-infections, emphasising the increasingly blurred lines between community and hospital-associated infections reported (Skov and Jensen, 2009).

In conclusion, we have presented the most detailed genomic analysis of MRSA isolated in Sri Lanka to date and have demonstrated, at least in the hospital and catchment area studied, that clinical MRSA infections in Sri Lanka are dominated by a PVL-positive ST5-MRSA-IVc clone. We have also shown the clone can be found in English patients with a history of travel to Sri Lanka. Further work is required to determine the prevalence of carriage and infection associated with PVL-positive ST5-MRSA-IVc in Sri Lanka, and the dynamics of transmission in and out of hospital, and whether these findings are replicated on a national scale.

## Author Contributions

TdS, AK and BP designed and supervised the study. SJS, EC, MU and SS carried out the field work and initial microbiological characterisation of the isolates. SC, SD and EB carried out data analysis and interpretation of the primary dataset. SM carried out genetic characterisation of isolates from LK and UK. BP performed phylogenetic analyses. SM, TdS, AK and BP performed analysis of study data. AK, BP, TdS and SM wrote the paper. GC, SP, CAA and LD provided data. All authors contributed to and approved the final manuscript.

## Acknowledgements

We thank the Staphylococcal Reference Service team at PHE, Colindale for their assistance with laboratory work. We also thank the microbiology laboratory staff at the Teaching Hospital Anuradhapura for help with collection and initial characterisation of isolates. We are grateful to Carlos M. Luna, Pulmonary Division, Department of Medicine, Jose de San Martin Hospital, University of Buenos Aires, Buenos Aires, Argentina, for providing bacterial isolates. This work was presented in part at the 26^th^ European Congress of Clinical Microbiology and Infectious Diseases, Amsterdam, The Netherlands.

## Access to data

The sequence data supporting the results of this article are available in the European Nucleotide Archive, under project accession number PRJEB27049.

